# Visualization and label-free quantification of microfluidic mixing using quantitative phase imaging

**DOI:** 10.1101/137117

**Authors:** GwangSik Park, Dongsik Han, GwangSu Kim, Seungwoo Shin, Kyoohyun Kim, Je-Kyun Park, YongKeun Park

## Abstract

Microfluidic mixing plays a key role in various fields, including biomedicine and chemical engineering. To date, although various approaches for imaging microfluidic mixing have been proposed, they provide only quantitative imaging capability and require for exogenous labeling agents. Quantitative phase imaging techniques, however, circumvent these problems and offer label-free quantitative information about concentration maps of microfluidic mixing. We present the quantitative phase imaging of microfluidic mixing in various types of PDMS microfluidic channels with different geometries; the feasibility of the present method was validated by comparing it with the results obtained by theoretical calculation based on Fick’s law.

## 1. Introduction

Microfluidic devices that control micrometre-sized samples in fluids are essential tools in various research fields such as analytical chemistry, biology and medicine [1, 2]. Mixing in microfluidic channels has great potential due to the capability of controlling droplet volume, chemical concentration, and sorting of fluids [3]. Recently, microfluidic mixing has been used for pharmaceutical developments and medical diagnostics [4, 5].

In order to monitor microfluidic mixing, various optical imaging techniques have been proposed and used [6]. For example, phase contrast microscopy, differential interference contrast microscopy, and confocal microscopy have been employed to observe the mixing and extract information of fluid mixture. Among various methods, bright-field and fluorescence microscopy are well-established and readily available [7]. However, existing techniques have limitations in providing quantitative imaging information. Most fluids are transparent and thus do not provide enough imaging contrast. Although the use of exogenous fluorescent or dye molecules can enhance imaging contrast, these techniques provide only qualitative information on the local concentrations of fluids and also raise the issues of altering intrinsic fluid conditions.

This imaging constraint is unfortunate because when microfluidic mixing can be quantified, microfluidic devices have much to offer with their unique control capabilities, high flexibility, and avoidance of the use of large amounts of samples. Despite the many challenges, there is a strong motivation to provide quantitative imaging of microfluidic mixing, especially if that imaging can be achieved without introducing exogenous labeling agents. Refractive index (RI) variation in microfluidic mixing can be detected by various optical methods including schlieren microscopy, speckle photography, and surface plasmon resonance [8-14]. However, these techniques do not provide quantitative measurements of microfluidic concentration.

To overcome the limitations in the imaging of microfluidic mixing, quantitative phase imaging (QPI) techniques can be employed. QPI is based on interferometric microscopy; this process quantitatively and precisely measures the optical phase delay maps introduced by a sample. Recently, QPI has been widely used for the study of various cells and tissues [15, 16]. Recently, QPI techniques have been employed with microfluidic channels. However, the quantitative analysis of mixing processes in microfluidic devices has remained unexplored. Previous works have used microfluidic channels only to verify developed QPI imaging systems or the loading of biological samples [17-24].

With QPI techniques, fluid concentrations in microfluidic mixing can be precisely quantified without using exogenous labels. For a microfluidic channel with a known height, the RI map of a fluid in the channel can be retrieved from the measured optical phase delay image. Importantly, the RI of a solution can be directly translated into the concentration of the solute via the linear relationship between the concentrations and the RI values [25].

Here, we present the visualization of microfluidic mixing in channels and quantitatively analyze the concentration of solutions during the mixing processes. A QPI technique based on Mach–Zehnder interferometry was used to obtain the optical phase delay maps of mixing solutions with different RIs [26], from which the concentrations of the solutions were calculated. Using this method, the concentration profiles of solutions in T-shaped microfluidic channels can be measured at various positions along the propagating axis. The experimental results were also validated with theoretical calculations based on Fick’s law. In addition, we quantified the mixing processes in microfluidic channels with complicated geometry, including three-input and contraction–expansion array (CEA) channels.

## 2. Methods

### A. Channel designs and conditions of fluids

Three types of polydimethylsiloxane (PDMS)-based microfluidic channels were used in this study: T-shaped, three-input, and CEA channels. The channels were fabricated by soft lithography. PDMS prepolymer and a curing agent (Sylgard 184; Dow Corning, MI) were mixed at a mass ratio of 10:1, and then cured for an hour on a hot plate at 75°C. The PDMS replica and a glass slide were treated with oxygen plasma (200 mTorr, 200 W) for bonding.

The T-shaped channel had a 9.4 μm height, 30 μm width of the input region, and 60 μm width of the mixing region. The three-input channel had 9.1 μm height, 30 μm width of the input region, and 90 μm width of the mixing region. The CEA channel had a 9.1 μm height, 30 μm width of contraction regions, 150 μm width of expansion regions, and 150 μm length of contraction and expansion regions.

Fluids used in the microfluidic channels were deionized (DI) water and sucrose solutions, which have different RI values. DI water and 20% (w/w) sucrose solution were used in the T-shaped and CEA channels. DI water at a level of a 20% and a 30% sucrose solution were used in the three-input channel demonstrations. A syringe pump (PHD ULTRA CP 4400, Harvard Apparatus, USA) was used to control the flow rates of all fluids at 1 μL/min.

### B. Optical setup

To obtain optical phase delay maps of fluids in microfluidic channels, Mach–Zehnder interferometry was used, as shown in Fig. 1. A coherent laser source (He-Ne laser, *λ* = 633 nm, 10 mW, Thorlabs Inc., USA) was used as illumination. The beam from the laser was first split into two arms by a beam splitter (BS1); the two beams were used as reference and sample beam, respectively. The diffracted beam from the microfluidic channel was collected by a long working distance objective lens (OL, LMPLFLN 20x, NA = 0.4, magnification 20×, Olympus Inc., Japan). The sample beam was further magnified four times using an additional 4-*f* configuration; the total magnification was 80×. The sample beam and the reference beam interfered at the camera plane; the interferogram was recorded by an sCMOS camera (C11440-22C, Hamamatsu K.K., Japan).

### C. Field retrieval algorithm

From one recorded hologram of the microfluidic channel, a complex optical field consisting of both the amplitude and the phase delay maps was retrieved by applying a field retrieval algorithm [Fig. 2] [27, 28]. A spatially modulated interferogram was first Fourier transformed, from which the modulated complex-amplitude field information was retrieved.

Amplitude and phase delay maps of the microfluidic channel are shown in Figs. 2(c) and (d), respectively, which were retrieved from the hologram in Fig. 2(b). Because the sucrose-water solution and the PDMS channels are transparent in the visible range, the amplitude images do not provide contrast [Fig. 2(c)]. On the other hand, because the RI of the sucrose-water solution depends on the solution concentration, light passing through solutions with different concentrations has a different optical phase delay [Fig. 2(b)]. The retrieved optical phase delay map shows a clear image contrast [Fig. 2(d)].

**Fig. 1.**
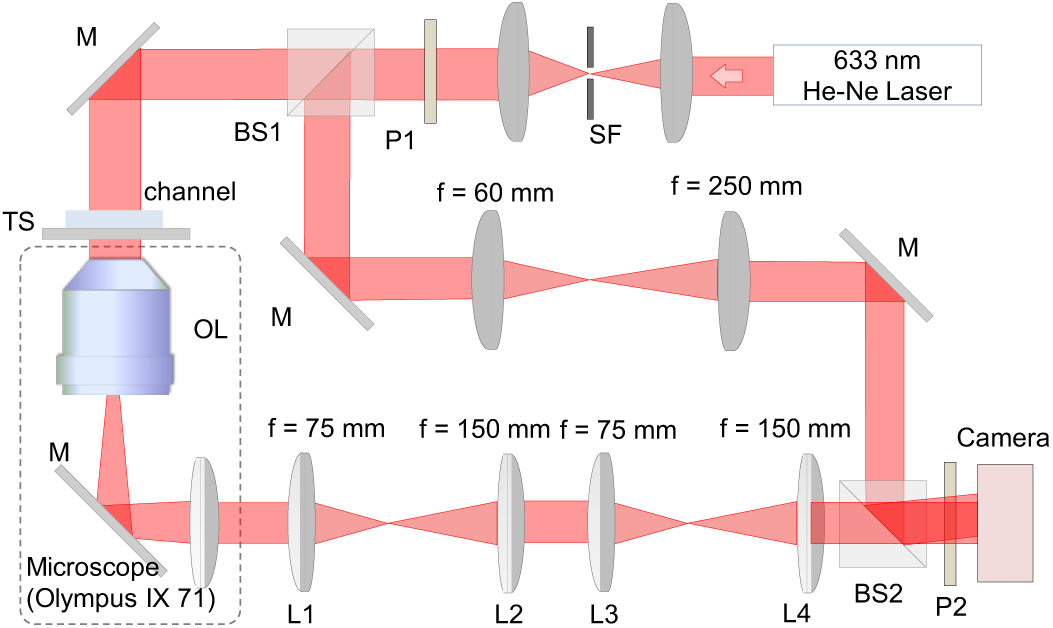
Quantitative phase imaging setup (SF: spatial filter, BS: beam splitter, M: mirror, TS: translation stage, OL: objective lens (Olympus 20×, NA 0.4), P: polarizer, L: lens).

**Fig. 2.**
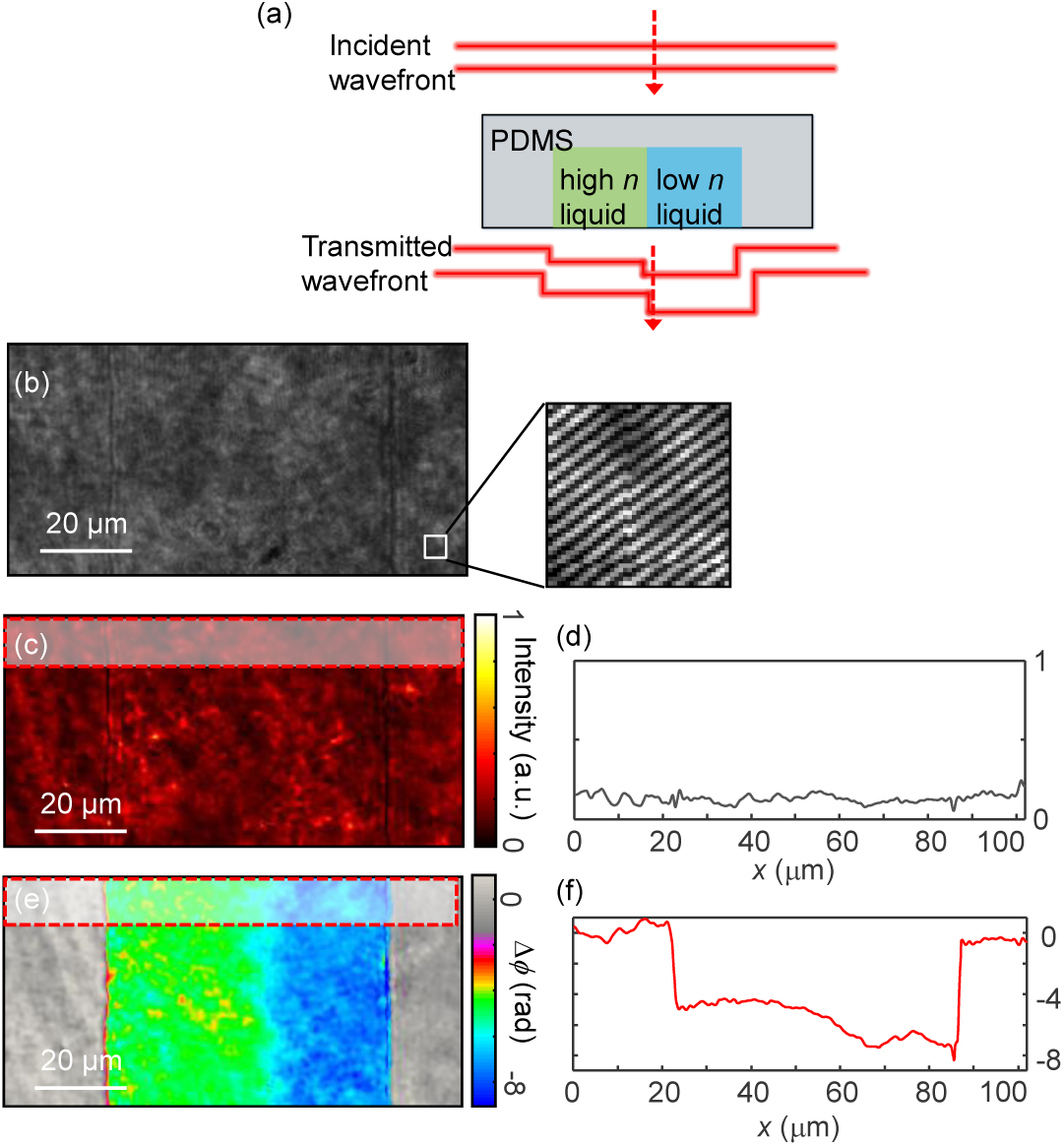
Representative optical phase delay map obtained from the microfluidic channel. (a) Concept of phase imaging, (b) a measured hologram of a channel, (c) amplitude, and (d) a phase image of the region in (b). Profiles of the red-dashed regions are shown as subplots after averaging along the flow directions.

To obtain a wide field-of-view (FOV) channel image, we obtained segmented optical field images at various positions of the microfluidic channel. A digital stitching method was used to connect these images into a large FOV image [26]. We employed a motorized translation stage (TS, ML203, Thorlabs Inc.) to translate the microfluidic channel in a fully automated manner. Each image overlapped 20% with adjacent images. The FOV of each segmented image is 166 μm × 166 μm, and the FOV of the stitched image extends to 700 μm × 1,000 μm. The acquisition time for obtaining each segmented image was 25 ms, and the total acquisition time was less than 45 s.

### D. Relation between RI and concentration of solutions

From the obtained optical phase delay map, the *Δϕ* of the PDMS microfluidic channel with known height *h*, and the RI value of the solution flowing inside the microfluidic channel can be calculated using the following equation, 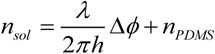, where *n*_sol_ and *n*_PDMS_ are the RI value of the solution and PDMS, respectively, and *λ* is the wavelength of the laser.

The RI of a solution is linearly proportional to its concentration. Thus, the mass concentration of a solution *c* can be described as: *c* = *c*_*b*_+ *β*Δ*n*, where *c*_*b*_is the mass concentration of the solvent, *Δn* is the RI difference between the solution and the solvent, and *β* is a proportional constant **[25]**.

### E. Theoretical analysis of mixing by diffusion

To validate the measurements, we compared the experimental results with the theoretical expectations based on Fick’s law [26-28]. We assumed a steady state flow condition and a constant flow velocity of the two fluids with different solute concentrations mixing in a T-shaped channel, where mixing occurs at the boundary of the two fluids. Fick’s second law of diffusion becomes a second order differential equation due to the given boundary conditions of the shape of the channel. As a result, the two-dimensional mass concentration distribution of fluids, c(x,y), flowing along the x-axis in the T-shaped channel of width w (y direction) can be described as:

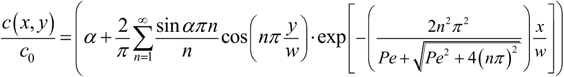

where *c*_0_ is the highest concentration, *α* is the relative position of the boundary of the two fluids in the input region ranging from 0 to 1 which can be determined based on the mass conservation, and *Pe* is the Pécket number, which describes the ratio of advective to diffusive transport rates.

## 3. Results and discussion

### A. T-shaped channel

In order to demonstrate the capability of the present method for quantifying microfluidic mixing, we first measured two fluids with different RIs, DI water (*n* = 1.333 at *λ* = 633 nm) and 20% sucrose solution (*n* = 1.364 at *λ* = 633 nm), mixing in a T-shaped channel [Fig. 3]. DI water was injected from the downside of the channel, while 20% sucrose solution was injected from the upper side of the channel.

Optical phase images of the fluids mixing in the T-shaped channel are shown in Fig. 3. The two fluids were injected from opposite ends at the initial positions of the T-shaped channel. Then, the two fluids mixed as they flowed through the channel. Figure 3(a) provides a phase map of the mixing fluids in the entire region (0.5 mm x 8 mm) of the T-shaped channel. Figures 3(b-e) are magnified phase maps indicated in Fig. 3(a) with dashed boxes; these maps depict locations 100 μm, 2100 μm, and 6100 μm from the starting point of the two-fluid mixing.

The measured optical phase images show a high image contrast due to the difference of the value of RI between fluids. Mixing of fluids is also clearly shown as they flow through the channel. At the input regions [Figs. 3(b)–(c)], a distinct boundary between the two fluids can be clearly seen, with a phase difference of 2.9 rad; this value agrees well with the theoretical expectation of 2.892 rad. After the liquids traveling a long distance along the channel, the boundary between the two fluids gradually disappeared, indicating the mixing of the fluids.

For quantitative analysis, optical phase profiles are extracted from the measured phase at various positions; these profiles are used to calculate the fluid concentrations [Fig. 3(f)]. Phase profiles at various positions along the y-axis were calculated by averaging values along the *x*-axis in each dashed box in Figs. 3(c)–(e). The profiles of the fluid concentration at various positions clearly and quantitatively show the mixing of fluids. The step-wise concentration profile at the initial position gradually changed into a concentration profile with a decreased concentration gradient.

To verify the measured results, we also compared the retrieved concentration profiles with the theoretical expectations calculated using Fick’s law, with results indicated by the solid lines in Fig. 3(f). The experimental and theoretical data exhibit good agreement.

### B. Three-input channel

To further demonstrate the capability of the present method, fluid mixing in a three-input channel was also measured. We used three fluids with different RIs: DI water (*n* = 1.333 at *λ* = 633 nm); 20% sucrose solution (*n* = 1.364 at *λ* = 633 nm); and 30% sucrose solution (*n* = 1.381 at *λ* = 633 nm).

Figure 4 shows quantitative phase images of fluid mixing in the three-input channel. A schematic of the channel geometry is provided in Fig. 4(a). DI water at a level of 20%, and 30% sucrose solution were injected from each inlet. Two distinct boundaries between fluids can be clearly seen in Fig. 4(b); the boundaries faded out as the fluids traveled along the channel.

**Fig. 3.**
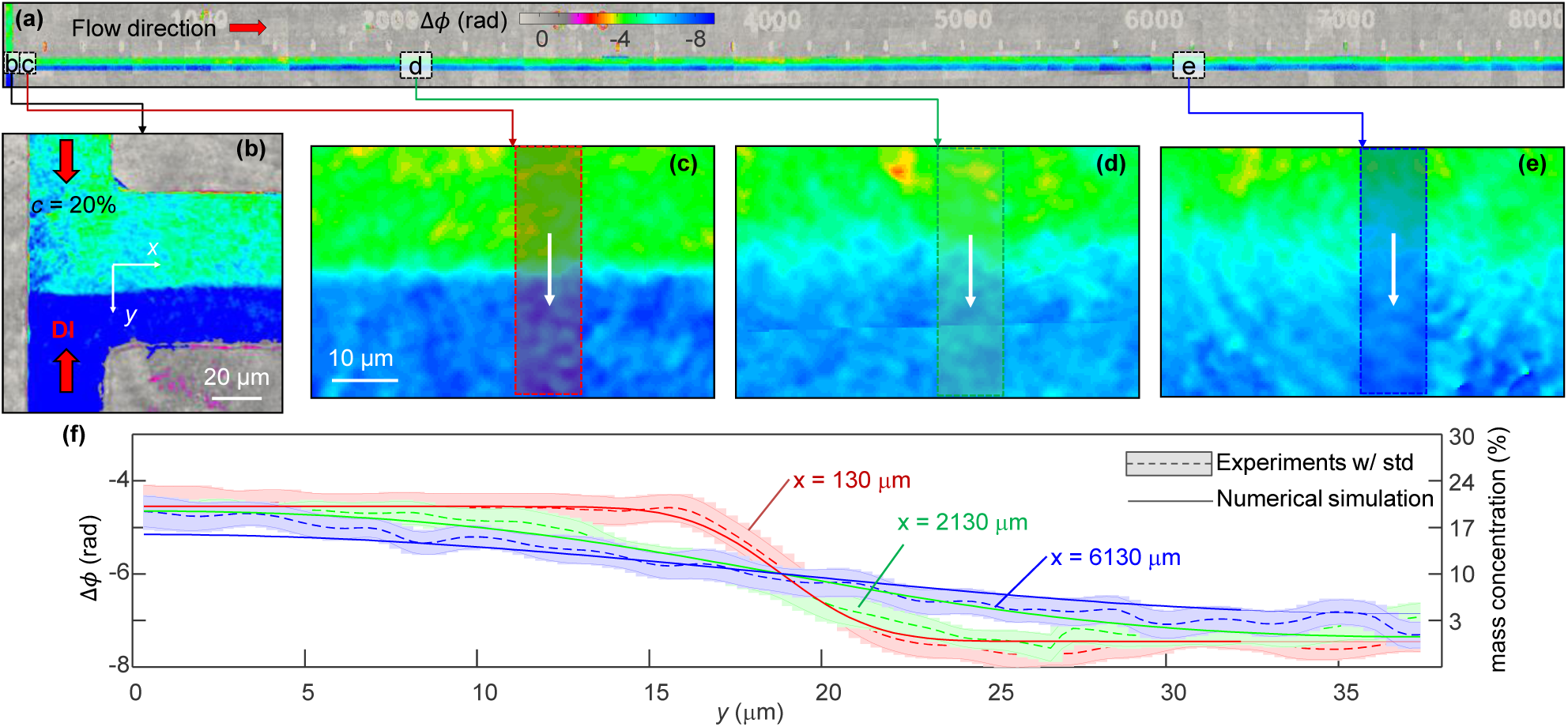
T channel mixing phase image (DI water and 20% sucrose solution mixing), (a) long image of T channel, (b-e) section images at each position, (b) image at input,(c) at 100–160 μm, (d) at 2,100–2,160 μm and (e) at 6,100–6,160 μm, (f) profiles of each dashed-section in arrow directions (red: mean (dashed line) at 130–140 μm and theoretical (solid line) values at 135 μm, green: mean (dashed line) at 2,130–2,240 μm and theoretical (solid line) values at 2,135 μm, blue: mean (dashed line) at 6,130–6,140 μm and theoretical (solid line) values at 6,135 μm).

**Fig. 4.**
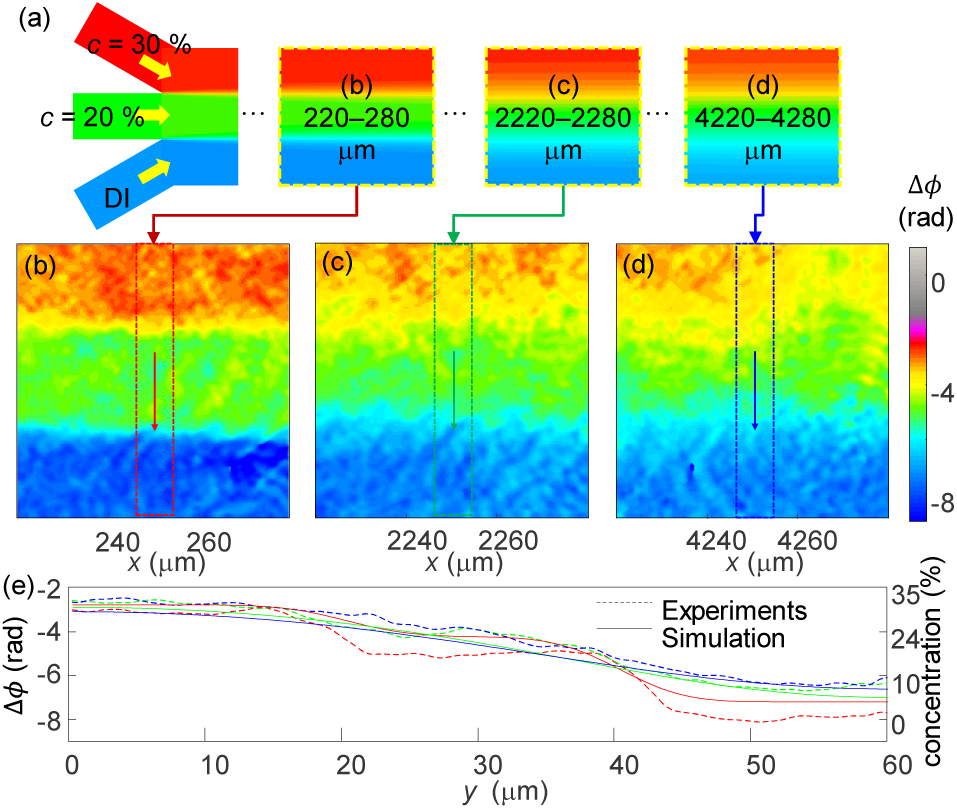
Three-input channel mixing phase images (mixing of DI water at 20% and 30% sucrose solutions), (a) three-input channel schematic image, (b) 200–300 μm, (c) 2,200–3,300 μm, (d) 4,200–4,300 μm regions and (e) theoretical (solid line) and mean (dashed line) profiles for each dashed box region in arrow directions.

At the initial position, the measured optical phase values are exactly consistent with the expected values. The relative phase differences at the boundaries of the neighboring solutions are calculated and found to be *Δϕ* = 2.800 rad between DI water and the 20% sucrose solution and 1.536 rad between the 20% and 30% sucrose solutions. The measured phase differences are 2.8653 and 1.9341, respectively. The measured phase images show the mixing of the three fluids as they travel. Figures 4(b)–(d) show phase images from different positions of the three-input channel 0.2 mm, 2.2 mm, and 4.2 mm from the initial position of the channel, respectively.

To quantitatively analyze the concentration distribution of the solutions during the mixing process, we extracted the phase profile from each phase image at different positions along the propagation direction and calculated the fluid concentration using the RI values of the solutions [Fig. 4(e)]. The measured profiles clearly show that the slopes at the boundary between two adjacent fluids are steeper near the entrances than at the back region.

We also calculated theoretical values of the concentration distribution based on Fick’s law, and compared these values with the experimental results. Although the overall shapes of the concentration distributions for the theoretical and experimental results matched, the exact concentration values exhibit some deviation. We assume that this mismatch can arise from the dilute solution assumption in Fick’s second law, which is derived based on the premise that the viscosities of the fluids are not significantly different. However, the solutions used have different viscosities, as can be seen in Fig. 4.

### C. CEA channel

To further demonstrate the applicability of the present method, we quantitatively measured the fluid mixing in the CEA channel. The CEA channel consisted of contraction and expansion regions with different widths; using the force arising from the channel shape, this type of channel has been used as an efficient mixing locus [29, 30]. When entering an expanding region with an abrupt change in the cross-sectional area, a centrifugal force is applied to fluids, which sequentially introduces Dean vortices and highly enhances mixing efficiency [31].

Figure 5 presents quantitative phase images of fluid mixing in a CEA channel. DI water and a 20% sucrose solution are injected into the input outlets, as depicted in the schematic [Fig. 5(a)]. The optical phase images of the third and ninth expansion regions inside the CEA channel are shown in Figs. 5(b) and (c), respectively.

**Fig. 5.**
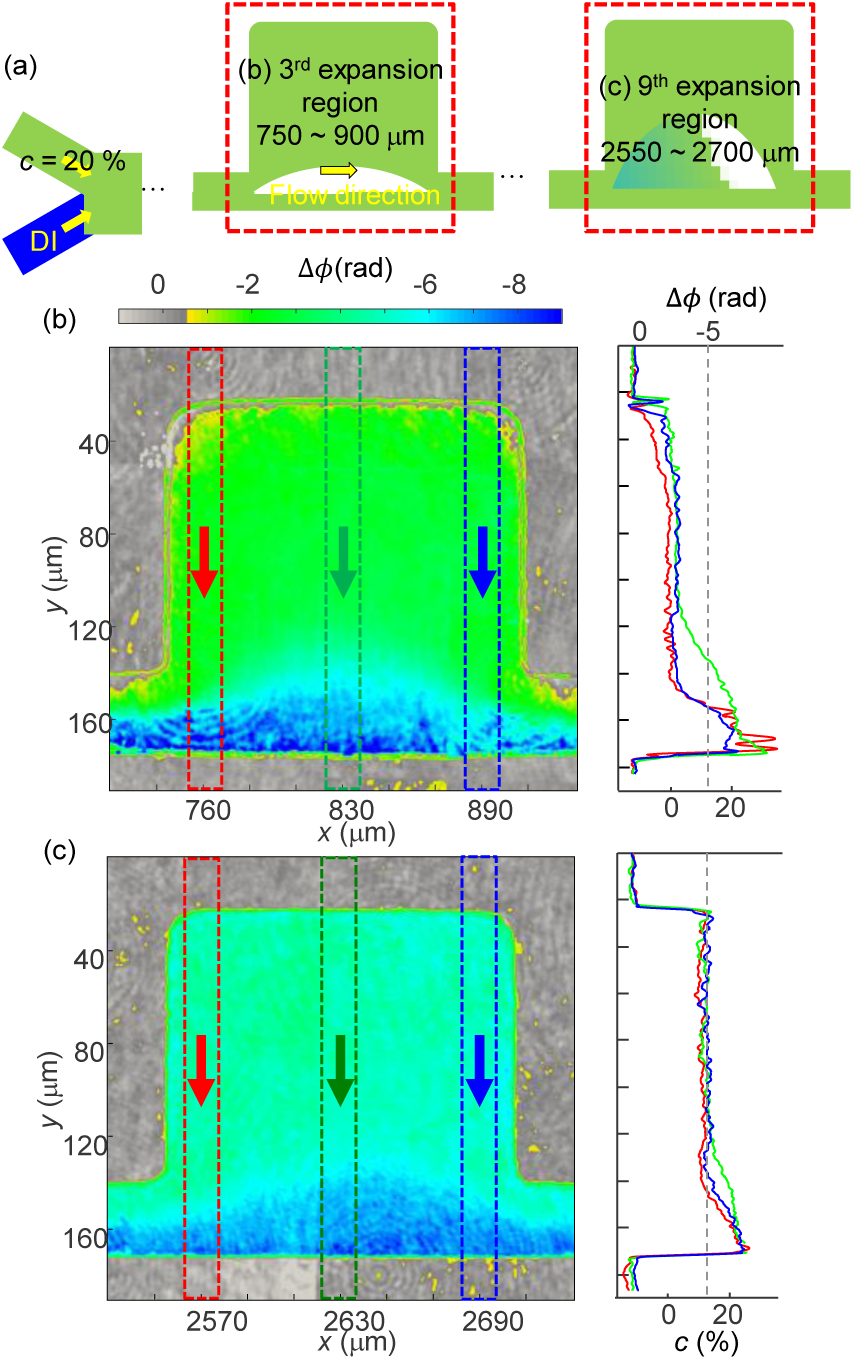
CEA channel phase images (DI water and 20% sucrose solution mixing), (a) CEA channel schematic image, phase image at (b) 3rd expansion region, (c) 9th expansion region. The right images in panels b and c indicate profiles (d) at 3rd and (e) 9th expansion region, respectively.

The optical phase delay maps clearly indicate the mixing of the fluids as they pass through the expansion regions. Compared to the image of the third expansion region, the phase image of the ninth expansion region shows decreased spatial gradients in the phase distributions, indicating the mixing of fluids.

To further analyze the measured results, we extracted the phase delay profiles at different positions along the propagation direction and calculated the concentration distribution. The three phase profiles from the front, center, and back of two expansion regions are averaged, as indicated by the dashed boxes in Figs. 5(b)–(c). The profiles at identical positions in each expansion region show that the slopes in the concentration distributions at the boundary between the two fluids are steeper near the entrances than they are at the exits.

## 4. Conclusions

In this work, we demonstrated a quantitative phase imaging of fluids mixing in microfluidic channels and a label-free visualization of concentration distributions in microfluidic mixing. The experimental results were validated in comparison with theoretical calculations based on Fick’s law. The applicability of the present method is also demonstrated with various microfluidic channels with different geometries.

This work presents the label-free quantitative imaging of microfluidic mixing, exploiting the RI values of various fluids. Although the quantitative phase imaging of microfluidic channels has been reported previously, quantitative imaging and analysis of fluid mixing has not yet been reported. Conventional approaches inevitably require the use of fluorescent or absorptive dyes; they also provide only qualitative information. However, the use of additional labeling agents is not necessary in the present method. Furthermore, the optical phase delay maps of microfluidic devices provide quantitative information about fluid concentrations.

Despite these advantages, the present method also has several limitations. First, quantitative phase microscopy or interferometric microscopy is required to measure the optical phase delay. But, these microscopes are relatively complicated and not widely available as compared to the devices used for conventional bright-field or fluorescence microscopy. Nonetheless, the recent advances in QPI provide several solutions: a QPI unit comparable to an existing microscope [32], or commercialized quantitative phase microscopes, are available [33]. The second limitation is that only fluids with distinct RI values can be visualized; fluids with similar RI values may be difficult to image with quantitative phase microscopes, which have a limited phase sensitivity, especially when speckle noise exists. However, this phase sensitivity issue can be remedied by exploiting low-coherent light, such as white light illumination or speckle illumination [34-37]. Third, different fluids with the same RI values cannot be distinguished with the present method because this approach does not provide molecular specificity. Nonetheless, spectroscopic phase measurements can be employed to some extent to address this issue of exploiting the optical dispersion [38-40].

Nonetheless, the label-free and quantitative imaging capability of the present method will find several important microfluidic applications in which the use of exogenous labeling agents for visualization causes limitations. Furthermore, measurements of the three-dimensional (3D) RI distributions of fluids can also be possible with optical diffraction tomography; this can further provide 3D concentration distribution information for mixing fluids without *a priori* information about the channel height [41, 42]. We envision that the use of RI will lead to various studies and experiments using microfluidic mixing.

## Funding

This work was supported by KAIST, Tomocube Inc., and the National Research Foundation of Korea (2015R1A3A2066550, 2014K1A3A1A09063027, 2014M3C1A3052567, 2016R1A2B3015986) and Innopolis foundation (A2015DD126). Prof. Park has a financial interest in Tomocube Inc., a company that commercializes optical diffraction tomography.

